# Amplicon sequencing detects, identifies, and quantifies minority variants in mixed-species infections of *Cryptosporidium* parasites

**DOI:** 10.1101/2025.04.10.648298

**Authors:** Randi Turner, Doaa Naguib, Elora Pierce, Alison Li, Matthew Valente, Travis C. Glenn, Benjamin M. Rosenthal, Jessica C. Kissinger, Asis Khan

## Abstract

*Cryptosporidium* is a globally endemic parasite genus with over 40 recognized species. While *C. hominis* and *C. parvum* are responsible for most human infections, human cases involving other species have also been reported. Furthermore, there is increasing evidence of simultaneous infections with multiple species. Therefore, we devised a new means to identify various species of *Cryptosporidium* in mixed infections by sequencing a 431 bp amplicon of the 18S rRNA gene encompassing two variable regions. Using the DADA2 pipeline, amplicons were first identified to a genus using the SILVA 132 reference database; then *Cryptosporidium* amplicons to a species using a custom database. This approach demonstrated sensitivity, successfully detecting and accurately identifying as little as 0.001 ng of *C. parvum* DNA in a complex stool background. Notably, we differentiated mixed infections and demonstrated the ability to identify potentially novel species of *Cryptosporidium* both *in situ* and *in vitro*. Using this method, we identified *Cryptosporidium parvum* in Egyptian rabbits with three samples showing minor mixed infections. By contrast, no mixed infections were detected in Egyptian children, who were primarily infected with *C. hominis*. Thus, this pipeline provides a sensitive tool for *Cryptosporidium* species-level identification, allowing for the detection and accurate identification of minor variants and mixed infections.

## Introduction

Species of *Cryptosporidium,* a genus of enteric protozoan pathogens, are leading causes of waterborne diarrheal diseases in humans and domesticated animals worldwide [1–4]. Both the Global Enteric Multicenter Study (GEMS) and the Study of Malnutrition and Enteric Diseases (MAL-ED) showed that cryptosporidiosis is the major cause of morbidity and mortality in children under five globally, leading to an estimated 48,000 deaths annually [5–7]. Infection typically occurs by consuming contaminated food or water. In low- and middle-income countries, endemic exposure derives from poor sanitation and hygiene [8, 9]. *Cryptosporidium* also continues to pose a substantial public health challenge in affluent nations, causing water or foodborne outbreaks and direct transmission within daycare facilities, hospitals, and other institutional settings [10–12]. *Cryptosporidium* oocysts are highly resistant to the chlorine-based disinfectants commonly used in water treatment and poses a challenge for standard filtration and water management [13]. Therefore, high-income countries experience outbreaks from contaminated recreational and drinking water [14].

There are over 45 recognized species and more than 100 genotypes of *Cryptosporidium* [15, 16]. While nearly 20 species have been reported in humans, most infections are caused by *C. parvum* and *C. hominis* [16, 17]. Species identification typically employs the small ribosomal subunit (SSU, 18S rRNA) gene [18, 19]. This locus includes both highly conserved regions amenable to universal primer design and variable regions capable of distinguishing between species. Other loci such as actin, heat shock protein 70 (HSP70), and cell outer wall protein (COWP) have been utilized, but high sequence conservation impedes resolving closely related species using these markers [20, 21].

Although the transmission modes are different between *C. parvum* (zoonotic) and *C. hominis* (anthroponotic), these species are closely related, sharing approximately 95% DNA sequence identity. They also share high sequence conservation with other human-infecting species in the *C. parvum* cluster, including *C. meleagridis* and *C. cuniculus* [17]. This similarity can make species identification difficult, particularly when species identification is solely based on 18S rRNA genotyping using the Sanger sequencing method. Further compounding this, the targeted Sanger sequencing method may entirely miss low-abundance genotypes in mixed infections [22]. Co-infections with multiple species of *Cryptosporidium* commonly occur, especially in highly endemic regions [23–25]. Co-infections frequently occur in livestock. Mixed *C. parvum* infections were widely reported in Uganda [26]. Subsequently, Grinberg *et. al.* showed an unprecedented intra-host genetic diversity, using HSP70 and *gp60* typing of two *Cryptosporidium*-positive fecal specimens collected in New Zealand. Next Generation Sequencing (NGS) identified these mixtures in samples from which only one allele per locus had been identified using Sanger sequencing [27, 28].

Early *Cryptosporidium* genotyping relied on restriction fragment length polymorphism (PCR-RFLP) digestion of 18S rRNA, producing species-specific banding patterns [29–31]. This time-consuming method often required additional sequencing steps to distinguish among closely related species. Other methods utilize 18S rRNA amplification in conjunction with Sanger sequencing or qPCR assays targeted at species-specific genes or regions within the 18S rRNA gene [15, 20]. These methods only poorly resolve mixed infections of closely related species.

NGS platforms generate a vast number of DNA sequence reads from any given sequencing reaction, allowing greater means to identify mixtures when they occur, and facilitate high-throughput parallelization of sequencing [32–34]. As a result, NGS provides means to accurately estimate the genetic diversity of *Cryptosporidium* species within a host. Here, we propose an amplicon sequencing approach targeting the V3/V4 region of the 18S rRNA gene. We utilize the DADA2 analysis pipeline in conjunction with a custom, curated *Cryptosporidium* 18S database for comprehensive species identification [35]. We demonstrate this approach’s sensitivity and its ability to accurately identify a broad range of species, to identify novel species and characterize the relative abundance of mixture constituents. We then demonstrate the application of this approach on veterinary and human samples, illustrating distinct patterns of transmission thereby resolved.

## Methods

### Ethics approval

This study was approved by the Research Ethics Committee of the Faculty of Veterinary Medicine, Mansoura University, with the code number MU-ACUC (VM.R.24.05.166). This survey followed the International Ethical Guidelines for Biological Research Involving Human Subjects.

### Sample collection

#### Rabbit stool sample collection

Between November 2022 and April 2023, a total of 230 fresh fecal samples were collected from 12 rabbit farms randomly selected in the Dakahlyia governorate in Northern Egypt. These small farms each housed 300-600 rabbits. Fecal samples were randomly collected from at least 25% of the rabbit cages on each farm, each sample consisting of 5-7 fresh fecal pellets from a given cage housing 4-6 rabbits. Fresh fecal pellets were then placed in sterile disposable plastic bags labeled with the age and breed of the animals (Flander, Baladi, Rex, Chinchilla, New Zeeland, and Hi-Plus), as well as the date of sample collection and location. At the time of sample collection, the animals appeared to be healthy. The rabbits ranged in age from 2 to 12 months and were divided into three age groups: ≤ 3-6 months old, >6–9 months old, and >9-12 months old.

#### Clinical stool sample collection

From May to August 2023, 330 stool samples were gathered from children in Northern Egypt experiencing diarrhea. Each person contributed one sample. The samples were obtained from four governorates: Dakahliya (n = 185), Damietta (n = 24), Gharbia (n = 71), and Sharqia (n = 50), reflecting regional variation in volunteer participation. These governorates were chosen for their proximity to Mansoura University to facilitate sample collection and storage. Before participating in the study, the parents or guardians of the children provided informed consent after being fully briefed on the study’s objectives. Once they consented, each parent or guardian completed a comprehensive questionnaire to gather epidemiological data, including gender, age, source of stool sample (hospital or clinic laboratory), contact with animals, and residency.

In a sterile tube, approximately 2 grams of fresh stool from each patient were mixed with 2 mL of 2.5% potassium dichromate. The mixture was homogenized thoroughly before storage at 4°C. Subsequently, samples were shipped for further processing to the Animal Parasitic Diseases Laboratory in Maryland, USA (Permit 20230214-0695A).

### Molecular Detection of *Cryptosporidium*

Total genomic DNA was isolated from 200 mg of stool using the DNeasy Powersoil Pro Kit (Qiagen, Hilden, Germany) according to the manufacturer’s instructions. Extracted total genomic DNA was screened for *Cryptosporidium* using real-time qPCR targeting the 18S rRNA gene using modified primers and probe set from Stroup *et. al* (GEMS Crypto 18S; Supplemental Table 1) [36]. Amplifications were carried out with 2 µL of extracted DNA in a total volume of 20 µL using a QuantStudio Flex 7. Reaction setup and cycling conditions can be found in Supplemental Methods. A red channel (Cyanine 5) collected data during the annealing/extension phase. Auto threshold function was utilized to calculate the threshold cycles (Ct) for each reaction using QuantStudio Real-Time PCR software program, (version 1.7.2). Samples were considered positive if their Ct value was less than 40. Positive samples were tested in triplicate.

### Primer Design for 18S *Cryptosporidium* Species Identification

Positive samples were selected for *Cryptosporidium* species identification by 18S amplicon sequencing. Forward and reverse primers were designed to target a 431 base variable region that spans the V3 and V4 regions (ILU Crypto 18S; Supplemental Table 1). Primers were modified to make them compatible with the iTru Adapterama indexes (ILU iTru Crypto 18S; Supplemental Table 1)

### *Cryptosporidium* 18S Database

To identify *Cryptosporidium* species, we created a custom *Cryptosporidium* 18S reference data set. We curated the available 18S rRNA sequences on CryptoDB (https://cryptodb.org/) [37, 38] and added other *Cryptosporidium* species sequences from NCBI (https://www.ncbi.nlm.nih.gov/) to expand species representation. The resulting custom reference data set consisted of 110 18S rRNA sequences, representing all 44 recognized *Cryptosporidium* species and various genotypes and environmental samples. The database can be found at https://github.com/jkissing/Turner_18S_Amplicon. Details on database curation can be found in the Supplemental Methods section.

### Species Identification and Abundance Estimation with DADA2

Illumina reads were processed to identify *Cryptosporidium* species and estimate their relative abundance using DADA2 (v1.26.0) in R (v4.2.2) [35]. Two rounds of taxonomy assignment were performed, beginning with genus determination by reference to the SILVA 132 NR99 SSU reference data set with a minimum bootstrap value of 50 [39]. For species of *Cryptosporidium,* we then performed a second round by reference to our custom 18S rRNA reference data set of known *Cryptosporidium* sequences, requiring a minimum bootstrap value of 55. To accommodate the possibility of a sequence representing a novel or highly divergent *Cryptosporidium* absent from the reference data set, we designated sequences assigned to the *Cryptosporidium* genus by reference to the SILVA database but lacking an exact match in our custom database as *“Cryptosporidium* sp.”. Sequences unclassifiable at any taxonomic level were labeled as “unassigned”. Details on read processing and abundance estimation can be found in the supplemental Methods.

### *In silico* Simulated *Cryptosporidium* Infections

Representative 18S rRNA sequences from various *Cryptosporidium* species were selected from NCBI. A “novel” *Cryptosporidium* species was created by intersplicing two variable regions from *C. andersoni* (OQ001483) into a *C. parvum* (AF112572) backbone (Supplemental Figure 3). For mixed infections, simulated reads from each species were pooled at defined frequencies (25%, 33% 50%, or 75%) to generate mock communities in the abundance file. Illumina reads were simulated using InSilicoSeq (v 2.0.1)[40] and processed in the DADA2 pipeline. Details on representative sequences and InSilicoSeq settings can be found in the Supplemental Methods.

### *Cryptosporidium* DNA Sources

*Cryptosporidium parvum* DNA was isolated from purified oocysts (Sterling Parasitology Laboratory, University of Arizona, Tucson, AZ). DNA from *C. hominis* (NR-2520) and *C. meleagridis* (NR-2521) was obtained from BEI Resources (https://www.beiresources.org/). DNA was amplified using the Repli-G Mini Kit (Qiagen, Hilden, Germany). Details on DNA isolation and amplification can be found in Supplemental Methods.

### Dilution Assay and Mock Infections

For the dilution assay, 1 µL of *C. parvum* DNA, ranging in concentration from 10 to 0.001 ng/µL, was spiked into 1 µL of extracted DNA from a rabbit stool sample confirmed as negative for *Cryptosporidium* by qPCR.

Single-species mock infections were created using 2 ng of DNA from *C. parvum, C. hominis,* or *C. meleagridis* DNA. Also, a total of 2 ng *C. parvum*, *C. hominis*, and *C. meleagridis* DNA were combined in prescribed compositions to create mock mixed infections.

### Illumina Sequencing of 18S Amplicon

18S amplicons were generated using Illumina iTru Crypto 18S primers (Supplemental Table 1) [41]. Amplifications were carried out with 5 µL extracted DNA and 2x Platinum II Hot-Start PCR master max (Invitrogen, Massachusetts, USA) according to manufacturer’s instructions. Amplicons were then barcoded using Adapterama I iTru indexes [42]. Samples were then pooled equally into a library for a final concentration of 4 nM and then prepared for sequencing using either Illumina V2 500 cycle Reagent kit or Illumina V3 600 cycle Reagent kit (Illumina, San Diego, CA), following manufacture guidelines. Amplicons were sequenced using a MiSeq, and the resulting reads were analyzed using the DADA2 pipeline. Additional details can be found in the Supplemental Methods.

### Sanger Sequencing of *Cryptosporidium* positive samples

Additional sample characterization was achieved by Sanger sequencing of amplicons from qPCR assays for 18S rDNA using external primers developed by Xiao *et al* (Xiao Crypto 18S external; Supplemental Table 1) [43]. BigDye cycle sequencing in each direction employed internal 18S primers (Xiao Crypto 18S internal; Supplemental Table 1) at a commercial sequencing service (Psomagen, Maryland, USA).

### Hierarchical Clustering

Forward and reverse reads were merged to create a consensus sequence using Geneious Prime 2020.2.4 software (Biomatters Ltd., Auckland, New Zealand; https://www.geneious.com/). Sequences were aligned using Clustal X/W and hierarchical clustering was achieved using Neighbor-Joining Geneious Prime (2024.0.2) with 1000 bootstrap replicates [44]. The consensus tree was generated with a midpoint root and 50% majority rule.

## Results

Developing amplicon sequencing for species-level identification of *Cryptosporidium* species.

We leveraged known variation in the 18S rRNA gene, the most widely used marker for *Cryptosporidium* species identification [1], to improve means to detect and estimate the relative abundance of constituents in mixed species infections. We sought a method that takes less time and achieves better characterization of mixed infections [23–25]. To do so, we developed an amplicon sequencing approach targeting a 431-base region of the 18S rRNA gene which encompasses the V4 region. This area contains sufficient sequence diversity to distinguish recognized *Cryptosporidium* species while also fitting within the size constraints of Illumina sequencing (Supplemental Figure 1).

After designing sequencing primers for this amplicon, we performed NCBI Primer-BLAST analysis (https://www.ncbi.nlm.nih.gov/tools/primer-blast/) and discovered that they are identical to other pathogens (*Blastocystis*, *Toxoplasma, Neospora, Theileria, Babesia, Eimeria, Hammondia hammondi, Cystoisospora belli, Cyclospora,* and *Isospora*). These other protozoan parasites also impose significant costs on human and animal health, worldwide. Thus, the tools described here have broad potential application (Supplemental Figure 2).

### *Cryptosporidium* Species Identification Workflow

Our workflow extracts DNA from stool samples and then screens for *Cryptosporidium* by real-time qPCR (Figure 1A). We considered samples with a Ct value of less than 40 positive. These we subjected to the described procedures for identifying and estimating the relative frequency of each species of *Cryptosporidium*.

**Figure 1.**
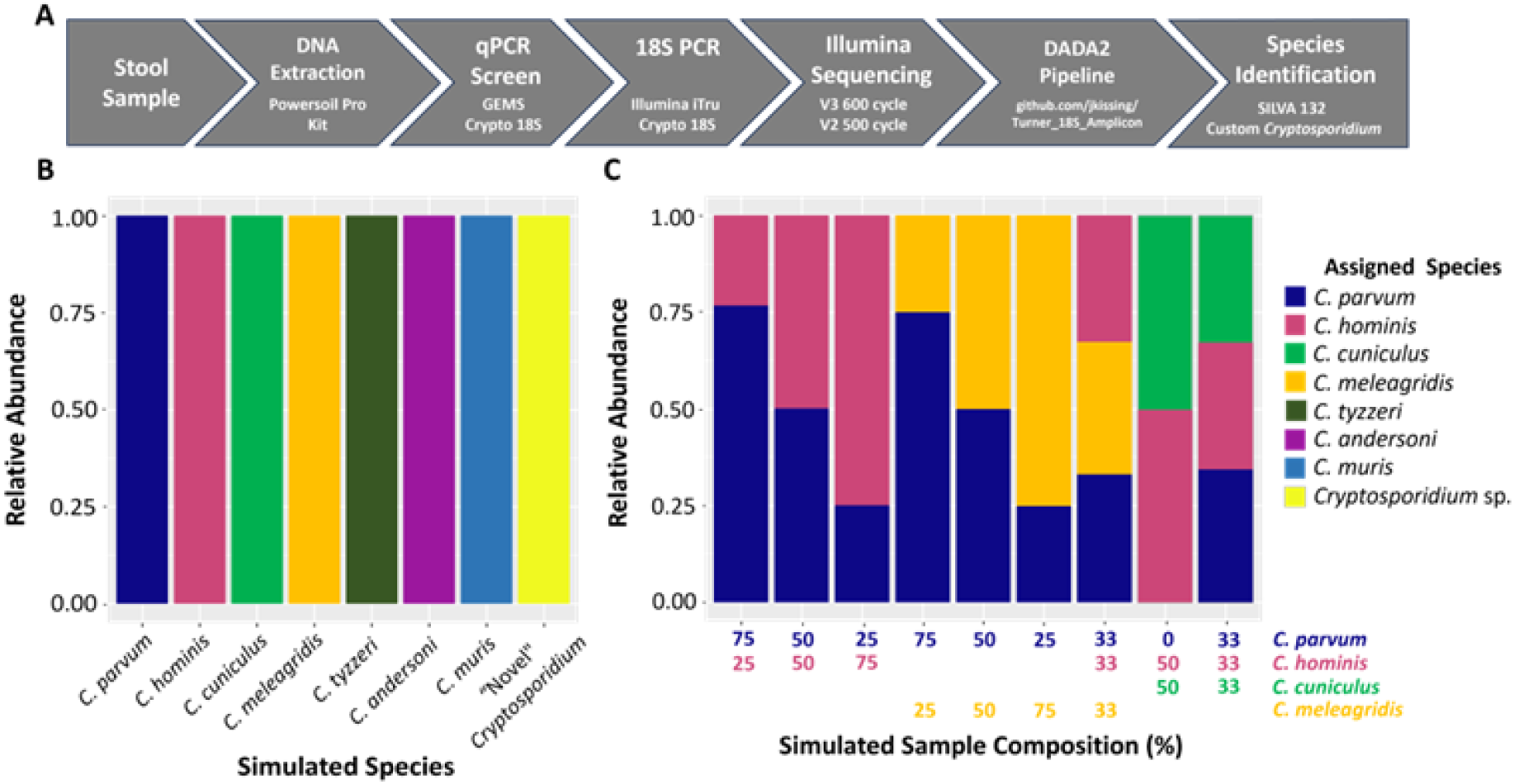
*Cryptosporidium* species identification using 18S amplicon sequencing. A) Schematic illustration of the amplicon sequencing and analyzing pipeline for species identification of *Cryptosporidium* from stool samples. DNA was extracted from stool and screened for *Cryptosporidium* by qPCR. 18S amplicons are generated from positive samples and sequenced using lllumina MiSeq. Reads were processed using the DADA2 pipeline. Taxonomy was determined using the DADA2 assign Taxonomy function in two steps: first at a genus level using the SILVA 132 reference database with a minimum bootstrap value of 50, then *Cryptosporidium* sequences were identified at a species level using a curated custom database and a minimum bootstrap value of 55. B) Bar diagram of relative abundance of identified species in simulated single-species infections. lllumina amplicon reads were simulated using lnSilicoSeq from selected *Cryptosporidium* species and a “never *Cryptosporidium* species made by intersplicing a variable region of C *andersoni* into C *parvum.* Simulated reads were mapped and processed using the DADA2 pipeline and taxonomy was assigned using the two-step taxonomy assignment. Each color indicates one single species. C) Relative abundance of species in simulated med infections. Iliumina am plicon reads were simulated form· ed-species infections in defined proportions using lnSilicoSeq Reads were then processed using the DADA2 pipeline, and species were identified using the two-step taxonomy assignment. Each color represents one single species. The absolute percentage of the contributed simulated reads of each species in them· ed infection is represented in the table at the bottom of the graph. Blank space indicates no contribution of simulated reads.

DADA2 uses a Naïve Bayesian Classifier algorithm to assign sequence taxonomy, referencing a broad data set [35]. Curating the contents of any such reference set lessens ambiguities or errors introduced by erroneous taxon assignments. Reference data sets also require curation to ensure adequate representation for the greatest interest.

Therefore, we first used the SILVA 132 SSU NR99 database to determine the genus associated with each sequence [39]. This database consists of over 690,000 sequences from all three major life branches, providing powerful means to distinguish among highly divergent organisms. However, it lacks sufficient detail to differentiate among all species of *Cryptosporidium*, as only representative sequences from *C. parvum* are present.

Therefore, we created a curated *Cryptosporidium* 18S rRNA database from sequences available at CryptoDB (https://www.cryptodb.org) [37, 38], adding selected sequences from NCBI (https://www.ncbi.nlm.nih.gov/). We removed sequences with ambiguous species origin based on BLAST and hierarchical clustering, resulting in a database consisting of 110 sequences representing 44 recognized *Cryptosporidium* species and a wide variety of genotypes and environmental samples (https://github.com/jkissing/Turner_18S_Amplicon). We performed a subsequent taxonomy assignment using this custom database to determine the *Cryptosporidium* species present.

### *In silico* Simulated *Cryptosporidium* Infections

We tested our identification pipeline by simulating both single-species and mixed infections of *Cryptosporidium in silico*. Illumina reads of the 18S rRNA gene from *C. cuniculus, C. hominis, C. parvum, C. meleagridis, C. muris,* and *C. andersoni* were subjected to our DADA2 identification pipeline after being simulated using InSilicoSeq [40]. When the simulated reads were mapped to both the SILVA 132 NR99 SSU reference dataset and our curated *Cryptosporidium* 18S rRNA reference database, all reads for each sample aligned with the corresponding 18S rRNA sequences of each tested species, demonstrating accurate identification of the simulated species across all test cases (Figure 1B). Importantly, the pipeline distinguished between closely related species, such as the most common species that infect humans (*C. parvum, C. hominis,* and *C. cuniculus*) which share > 95% DNA sequence identity (Figure 1B).

We then evaluated, *in silico,* our ability to estimate the composition of mixed infections employing 18S rRNA sequences from two or three human-infecting species in defined proportions (Figure 1C). The 18S rRNA-based amplicon sequencing method correctly identified all species simulated in a sample and accurately estimated the simulated abundance in every mixture (Figure 1C). To assess whether the method could identify a new species of *Cryptosporidium*, we simulated a “novel” *Cryptosporidium* species by interspersing variable regions of *C. andersoni* into a *C. parvum* backbone (Supplemental Figure 3). Our two-step identification method successfully classified the sequence as belonging to an unnamed species in the genus *Cryptosporidium* (Figure 1B).

### *In vitro* Mock Infections

To further assess our 18S rRNA-based detection system, we created *in vitro* mock infections. These included single species “infections” (adding 2 ng *C. parvum, C. hominis,* or *C. meleagridis* gDNA to rabbit stool). All ensuing reads were accurately assigned to the known species (Figure 2A). When rabbit stool was seeded with DNA mixtures, the process identified only those constituents as present (Figure 2C), in proportions that resembled (but did not perfectly reproduce) their target proportions. We discuss an apparent bias towards *C. parvum* below.

**Figure 2.**
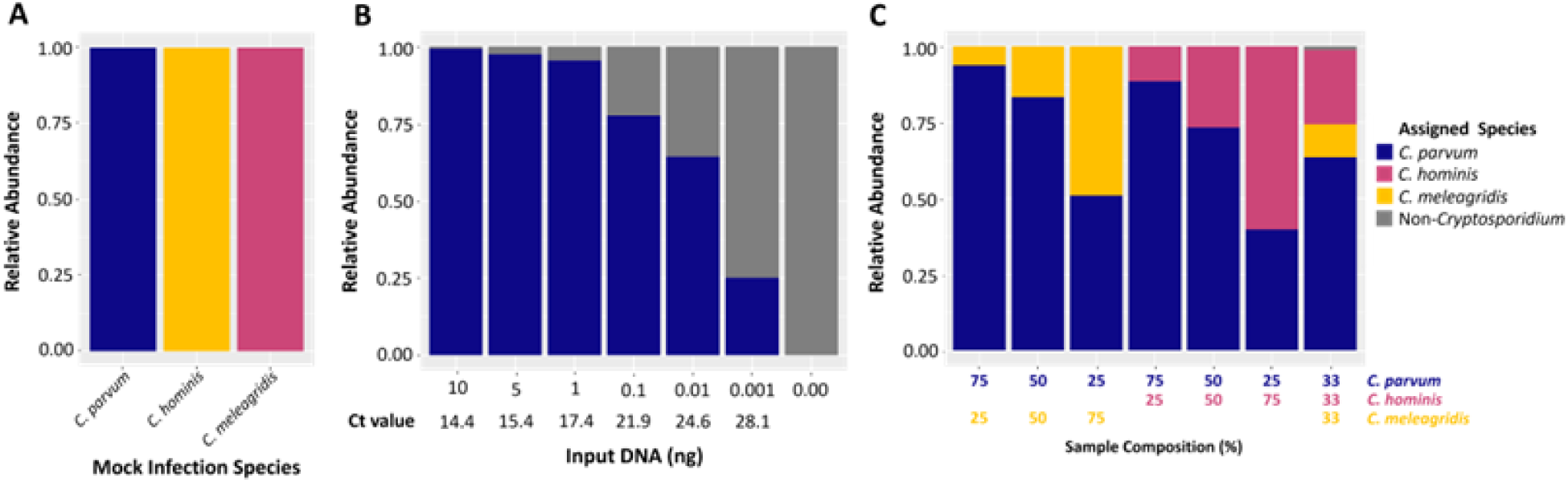
Estimation of sensitivity and specificity of 185-based amplicon sequencing and the calculation of relative abundance of *Cryptosporidium* species identified in mock infections. A) The specificity of *Cryptosporidium* detection using three different species. The specificity of the amplicon sequencing to identify different Cryptosporidium species is depicted in the relative abundance bar plots after assigning short reads of the 185 gene of C. parvum, C. hominis, and C. meleagridis in mock infections. B) 1sitivity of *Cryptosporidium* detection in a complex stool background. Varying amounts of *C paNum* DNA were diluted with DNA extracted from rabbit stool that was negative for *Cryptosporidium.* Sam pies were then amplified for 185 using ITru Crypto 18S primers, sequenced, and species identifications were performed using our DADA2 pipeline. The Y-axis indicates the relative abundance whereas the X-axis indicates the amount of C. parvum gDNA (ng) pooled into rabbit gDNA. CT value represents the qPCR value corresponding to each dilution. C) Precious detection of m· ed infections of three closely related *Cryptosporidium* species in mock experiments. A total of 2 ng C. vum, *C. hominis,* and *C. meleagridis* DNA were combined in prescribed compositions to create mock infections. Samples were then amplified for l&S using ITru Crypto 185 primers, sequenced, and species identifications were performed using our DADA2 pipeline. The Y-axis indicates the relative abundance of short-reads sequenced from each species. The X-axis indicates the prescribed composition to create each mock infection. The percentage of each species’ contribution to creating each mock mixed infection is depicted at the bottom of the figure.

### *In vitro* Sensitivity Assessment of *Cryptosporidium* Detection

To test the sensitivity of our approach, we spiked between 10 and 0.001 ng of *C. parvum* DNA into DNA from a complex stool background (Figure 2B). All or nearly all such reads attributed to *Cryptosporidium* were correctly assigned to *C. parvum* providing one measure of process validity. From stool samples containing no natural or added *Cryptosporidium,* no 18S reads were falsely ascribed to the genus (providing further assurance of method specificity).

Using qPCR standardized in The Global Enteric Multicenter Study (GEMS) study [5–7], we found that a C_T_ value of 28.1 (corresponding to 0.001 ng of *C. parvum* gDNA) was optimal for detecting constituents when present at a relative abundance of 25%. Interestingly, the proportion of non-*Cryptosporidium* reads decreased significantly as *C. parvum* input gDNA amount increases.

### In vitro mock mixed infection

To determine whether the method can effectively differentiate mixed infections and calculate the relative abundance of each species in a sample, we created artificial mixtures by combining 2 ng of gDNA from *C. parvum*, *C. hominis*, and *C. meleagridis* in defined amounts. The abundance estimations of the mixed infections revealed an overrepresentation of *C. parvum* in all mixtures (Figure 2C). This was particularly noticeable in mixed infections with *C. meleagridis*, where the method over estimated *C. parvum* abundance by up to two-fold (compared to the input concentration we attempted to add) (Figure 2C). We therefore sought to attribute such discrepancies to preferential amplification of *C. parvum* or to error in estimating gDNA concentrations added to the mixtures. To investigate this further, we assessed the CT values from qPCR using 2 ng of total gDNA from *C. parvum, C. hominis,* and *C. meleagridis*. The qPCR results confirmed that we had added more *C. parvum* to the mixtures than we had thought. The respective CT values for *C. parvum, C. hominis,* and *C. meleagridis* were 14.36, 15.24, and 15.56, respectively. This suggests that *C. parvum* enjoyed a two-fold concentration advantage over *C. meleagridis,* further validating read abundance as an indicator of true mixture composition.

### Prevalence of *Cryptosporidium* in Egyptian Rabbits

We tested our entire pipeline using rabbit stool samples from the Dakahalia governorate in northern Egypt. Previous studies estimated that 11.9% of this region’s rabbits harbor infections with *C. cuniculus* [45]. We collected 224 rabbit stool samples from 12 different farms; animal age ranged from 2 to 12 months. Total genomic DNA was extracted from the stool samples and screened for *Cryptosporidium* using quantitative PCR (qPCR). Of these, 17 samples tested positive by qPCR, exhibiting high C_T_ values (>35) suggesting low parasitemia. Repeat testing did not yield consistent results, likely owing to low target abundance (Supplemental Table 2). Of these 17 presumptively positive samples, our tool identified *Cryptosporidium* in 8. In 5 of these samples, *Cryptosporidium* reads accounted for fewer than 2% of all 18S reads; most of these were ascribed to *C. parvum* (Figure 3B). Additionally, *C. hominis*, *C. meleagridis,* and an undetermined *Cryptosporidium* sp. were detected three samples (R034, R186, and R195). From these rabbit fecal samples we detected reads from unspecified species in R154, R186, R197, and R198 (black bars, Figure 3B).

**Figure 3.**
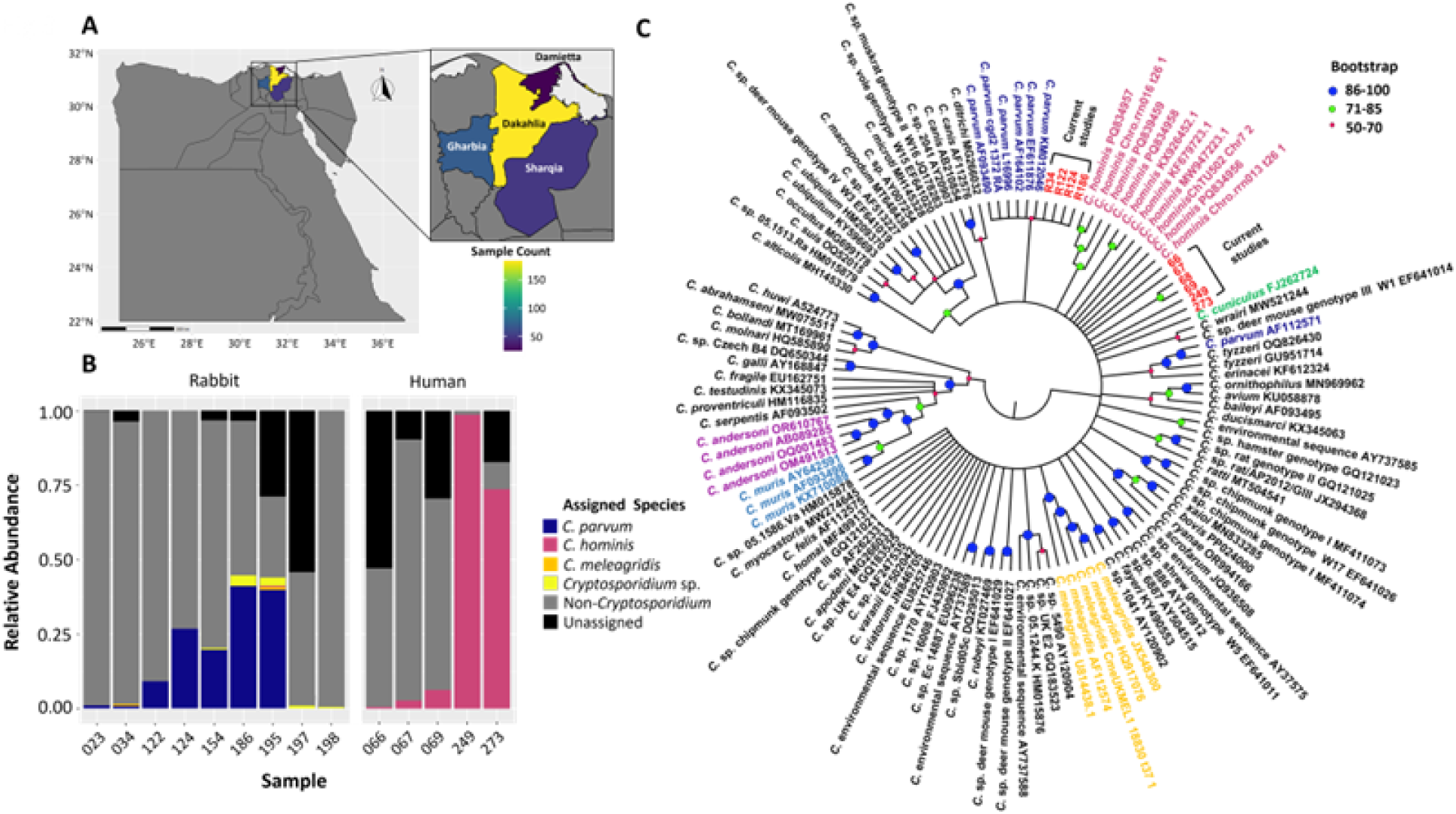
Detection and identification of *Cryptosporidium* species present in rabbit and clinical samples. A) Map of Egypt with the sample origin governorates colored by number of clinical samples from each govern ate. B) Relative abundance of *Cryptosporidium* species identified in rabbit and human clinical samples from Egypt. Samples were processed cording to the pipeline shown in Figure lA. Briefly, stool samples were amplified for 18S using ITru Crypto 185 primers and sequenced using lllumina MiSeq. Species identifications were performed utilizing our DADA2 pipeline with SILVA 132 reference <u>database,</u> followed by the curated custom *Cryptosporidium* <u>database</u> as mentioned in Jre **1.** Each color indicates each species, except black color, representing unassigned, and gray color, representing *non-Cryptosporidium* species. Each bar represents the relative distribution of *Cryptosporidium* species in each sample. C) Confirmation of 18S amplicon sequencing-based species identification with conventional genotyping methodology using Sanger sequencing of an amplicon amplified and sequenced by Crypto 18S primer sets. A Neighbor-Joining phylogenetic tree was constructed using ltol https://itolembl.de/itol.cgi) with 1000 bootstrap replicates after aligning the sequences using Cluster X/W. *Cryptosporidium* species names followed by NCBI accession number https://www.ncbi.nlm.nih.gov/l are provided in the phylogenetic tree. Sequenced samples from the current study are depicted in red color. The circle size at each node indicates bootstrap values.

To confirm species identifications and to compare the sensitivity of the Illumina-based detection system, we amplified a ∼1300 base pair fragment of the 18S rRNA gene using well-established *Cryptosporidium* species identification primers developed by Xiao *et. al.* [43]. We successfully amplified fragments from only four samples (R034, R122, R124, and R186). Sanger sequences of these four samples clustered with *C. parvum* in an unrooted phylogenetic tree (Figure 3c), reinforcing the identifications made through Illumina sequencing. Notably, none of the samples exhibited ambiguous bases expected when Sanger sequencing is applied, directly without cloning, to amplifications derived from mixed templates. However, the methods introduced here identified two samples (R034 and R124) with faint signatures of mixed infections involving *C. meleagridis*, *C. hominis*, and unidentified *Cryptosporidium* species. Thus, our new methods appear more sensitive for identifying mixed infections of *Cryptosporidium*.

### Prevalence of *Cryptosporidium* in Egyptian Children

To understand whether our Illumina-based detection system can accurately identify the species of *Cryptosporidium* present in children, determine the role of zoonotic transmission, and accurately calculate the rate of mixed infections, we collected 330 clinical samples between May and August 2023 from four Egyptian governorates: Dakahliya, Damietta, Gharbia, and Sharqia, located in the Nile Delta (Figure 3B). The samples were taken from children aged between 6 months and 10 years.

From 330 samples derived from Egyptian children, our method identified 22 (6.6%) for one or more species of *Cryptosporidium.* 14 of these came from 221 hospitalized patients and 8 from 109 persons visiting clinics (Supplemental Table 3). We did not find any significant correlation of infections with the distribution of age, sex, or urban versus rural sampling regions (Supplemental Table 3). The oocyst burden in the clinical samples varied extensively, indicated by a wide range of CT values in qPCR from 16 to 38 cycles.

To identify the *Cryptosporidium* species, present in these clinical samples, we used two approaches: 1) conventional species detection utilizing Sanger sequencing with commonly used PCR primers, and 2) species-level detection using 18S rRNA-based amplicon sequencing. For the conventional detection via Sanger sequencing, we attempted to amplify the 18S rRNA gene using all 22 qPCR-positive samples (Supplemental Table 3, Figure 3B). Only six of these 22 samples yielded a positive amplicon for Sanger sequencing. This sequencing identified *C. hominis* in all such positive samples (Supplemental Table 3, Figure 3B). Five of these samples (and no other sample) tested positive using our new amplicon sequencing method. Amplicon-based Sanger sequencing confirmed *C. hominis*, in all these samples, with no indications of mixed infections (Supplemental Table 3, Figure 3B, 3C). These results reaffirm *C. hominis* as a major causative agent of cryptosporidiosis in Egyptian children.

## Discussion

Although recent developments in molecular detection and genotyping methodology identified many cryptic species of *Cryptosporidium*, existing genotyping tools fail to accurately assess the relative abundance of the constituents of mixed infections. Therefore, we developed a sensitive and specific 18S amplicon-based short-read sequencing pipeline, followed by a two-step species assignment using the SILVA SSU database and a curated *Cryptosporidium* 18S-rRNA gene database. We determined this process specific, sensitive, and accurate in estimating the contributions of even closely related species of *Cryptosporidium* comprising mixed infections. Further, our amplicon-based detection system proved capable of identifying novel species of *Cryptosporidium*.

We then applied this procedure to real veterinary and human fecal samples, estimating prevalence rates and species compositions in a sample of rabbits and a sample of children. Rabbit samples identified *C. parvum* and an undescribed species of *Cryptosporidium*. Four of 8 rabbit samples appeared to be comprised of mixed-species infections. We estimated *Cryptosporidium* in 6.6% of the sampled Egyptian children. Most reads derived from these children corresponded to *C. hominis*, comporting prior estimates derived from existing genotyping methods. The amplicon-based system afforded greater means to identify mixed infections and unclassifiable sequences, which deserve further study as possibly undescribed parasite species. Thus, amplicon sequencing enables new insights concerning the dynamics of cryptosporidiosis which should enhance disease control strategies employing targeted interventions.

The molecular detection of different *Cryptosporidium* species is still in its infancy, and a cost-effective diagnostic tool for *Cryptosporidium* that offer improved sensitivity, specificity, and determine mixed infection is urgently needed. In enhancing the sensitivity and reducing the assay time, quantitative PCR (qPCR) has become an integral tool for the screening of environmental pathogens [46, 47]. Recently, notable progress has been made in the validation of fluorescence in situ hybridization (FISH) probes for the identification of the human infectious *Cryptosporidium* species, specifically *C. parvum* and *C. hominis* [48]. Probes Cpar677 (targeting *C. parvum*) and Chrom253 (targeting *C. hominis*) were validated by Alagappan *et. al.* [49]. The specificity and cross-reactivity of these probes were assessed against only eight *Cryptosporidium* species, *including C. andersoni, C. muris, C. meleagridis,* and *C. felis*. Furthermore, through careful primer design and melting curve analysis, it has now been established that pathogenic *Cryptosporidium* species can be differentiated from non-pathogenic ones [46, 47]. An important illustration of this capability was demonstrated by Li *et. al.* [50], who employed fluorescence resonance energy transfer probes and melting curve analysis in the 18S-LC1 and *hsp90* assays to distinguish common human-pathogenic species such as *C. parvum, C. hominis, and C. meleagridis*. The 18S-LC2 assay was also effective in differentiating non-pathogenic species, such as *C. andersoni*, from pathogenic species frequently detected in source water [50]. Hence, to overcome these challenges and develop a methodology to not only identify the known species but also the relative abundance of mixed infection and identification of the presence of new species, we developed a highly sensitive (0.001ng target DNA, C_T_ = 28) 18S-based amplicon sequencing for *Cryptosporidium*.

Over the past decades, the emergence of readily accessible high-throughput sequencing of amplicons derived from the 18S rRNA gene has fundamentally transformed the fields of clinical and public health microbiology [51–53]. The advancement of *Cryptosporidium* detection using amplicon sequencing has not only accelerated the process of pathogen identification but has also enhanced the accuracy of these identifications. Moreover, the integration of high-throughput methodologies with sophisticated bioinformatics tools will allow researchers to gain deeper insights into various critical aspects of infectious diseases, including how they are transmitted and the role of mixed infection in disease susceptibility. This approach is particularly vital for evaluating the diversity of *Cryptosporidium* in different ecosystems.

Previously, metabarcoding assays enabled the successful detection of protozoan parasites including *Cryptosporidium* [53–55]. However, none of those methodologies provided the tools to identify all known species, particularly closely related and novel species of *Cryptosporidium*. Taxonomy can only be assigned if a representative sequence is present in the reference data set. However, it is also possible that a *Cryptosporidium* sequence could originate a novel, highly divergent, or unrepresented species that is not found in the reference data set and thus would not be assigned a species. To counteract this, we leveraged the genus assignment from the SILVA database so that any sequences identified as originating from the *Cryptosporidium* genus are considered unspecified *Cryptosporidium sp.* Our two-step detection system using the SILVA database, and a curated *Cryptosporidium* 18S rRNA-gene database provides the tools to detect all known *Cryptosporidium* species in addition to unknown novel species.

The hybridization of parasites has emerged as a pressing public health concern [56]. The phenomenon of coinfections, where multiple species of eukaryotic pathogens simultaneously infect a single host, can have profound implications not only for the parasites themselves but also for their hosts. These interactions can manifest in various ways, including synergistic effects that enhance the virulence or spread of infections, or antagonistic effects that may hinder the establishment and impact of one or both parasites [57]. Additionally, hybridization also has a wide range of effects on drug efficacy in parasites [58]. Thus, it is critical to determine the frequency of mixed infections, and the relative abundance of the mixed infected species present in animal and clinical samples. It has been documented that mixed infection of *Cryptosporidium* at the species level is common in animal hosts. Few reports are available on clinical samples, which could be due to multiple factors including host immune response and low resolution of the current Sanger sequencing-based detection systems.

Our amplicon sequencing method correctly assigned and determined the relative frequency of each species including close and distantly related species of *C. parvum* using *in silico* and mock experiments. A notable study that employed amplicon-based next-generation sequencing (NGS) to identify various *Cryptosporidium* species found that an alarming 30% of the infected animals studied had mixed infections. Detection of mixed infections was made possible through NGS as Illumina sequencing technology conducts sequencing of each strand with massively parallel capabilities [59].

*Cryptosporidium* species are widely distributed among humans and animals, exhibiting both anthroponotic and zoonotic transmission cycles in Egypt [60]. Several species infect humans and animals have been reported in the country, including *C. parvum, C. hominis, C. meleagridis, C. ryanae, C. andersoni, C. xiaoi, and C. bovis*. Among these, *C. parvum* is the most prevalent, followed by *C. hominis* in human infections in individuals under 10 years old [60]. To gain a deeper understanding of the epidemiology, including transmission dynamics and differences in clinical presentations, as well as the frequency of mixed infections, it is essential to conduct more extensive molecular typing studies in Egypt. In this context, we carried out a surveillance study involving clinical and rabbit samples. Our findings revealed a prevalence rate of 7.6% in rabbit samples and 6.6% in clinical samples, which aligns with previously published prevalence rates of *Cryptosporidium* in animals and clinical cases [45, 61–63].

The comparative analysis of sensitivity between qPCR and amplicon sequencing revealed that samples with a C_T_ value above 31 are generally negative in amplicon sequencing, limiting the number of samples entered into the amplicon-based species detection pipeline. This finding confirms that the real-time PCR assay has higher sensitivity compared to the metabarcoding assay. It is crucial to note that the accuracy of quantitative detection via qPCR is contingent on the reliability of the standard curve, which can be influenced by pipetting errors and potential DNA losses. Amplicon sequencing showed that the clinical samples were primarily infected with *C. hominis*, whereas *C. parvum* was the major species detected in rabbit samples. This finding was also supported by the conventional method of species detection with the Sanger sequencing of the 18S gene. Notably, amplicon-based sequencing identified trace amounts of short reads from other *Cryptosporidium* species in several rabbit samples alongside *C. parvum* reads, indicating the occurrence of mixed infections in Egyptian rabbits.

Taxonomic profiling of complex microbial communities using 16S rRNA marker gene surveys has garnered significant interest, providing valuable insights into the bacterial composition of these communities and their links to health and disease [64, 65]. To deepen our understanding of species diversity and the relative abundance of mixed infections in both animal and clinical samples, we developed a highly specific and sensitive stepwise protocol that employs amplicon sequencing of the hypervariable regions of the 18S rRNA gene. While this protocol allows us to determine species composition and the relative abundance of parasite burden in clinical samples, we will need high-resolution and sensitive genotyping tools, such as capture enrichment [66, 67] or single-cell sequencing [68], for a more comprehensive investigation. These advanced methods could prove instrumental in unraveling the genetic complexity of this parasite, thereby enhancing the diagnosis and understanding of the genetic basis of cryptosporidiosis. The insights obtained from these findings can be applied to refine parasite detection methodologies, ultimately strengthening efforts to control and prevent intestinal parasitic infections.

## Data Availability

Curated *Cryptosporidium* 18S database, DADA2 pipeline scripts, and simulated Illumina reads are available at https://github.com/jkissing/Turner_18S_Amplicon. Sequencing data for all mock infection and clinical samples are available in NCBI BioProject database under accession number PRJNA1197520. Sanger 18S sequences from the human and rabbit samples are available in GenBank under accession numbers PQ963461-PQ963469.

## Acknowledgements

We acknowledge CryptoDB (https://cryptodb.org/) [37, 38] for providing a publicly available repository for *Cryptosporidium* genomic data. CryptoDB is part of the Eukaryotic Pathogen, Vector, and Host Informatics Resources (VEuPathDB, https://veupathdb.org/), VEuPathDB receives funding from the Wellcome Trust (UK) to support informatics efforts focusing on kinetoplastida and fungal organisms with special emphasis on improving functional annotation of genomes. Additional computing resources were provided by the SciNet HPC Consortium (https://scinet.usda.gov/). This research was supported in part by an appointment of Randi Turner and Doaa Naguib to the Agricultural Research Service (ARS) Research Participation Program administered by the Oak Ridge Institute for Science and Education (ORISE) through an interagency agreement between the U.S. Department of Energy (DOE) and the U.S. Department of Agriculture (USDA). ORISE is managed by Oak Ridge Associated Universities (ORAU) under DOE contract number DE-SC 0014664.

## Author contributions

Conceptualization: A.K., R.T., methodology: A.K., R.T., D.N., E.P., A.L., M.V.; software: A.K., R.T.; analysis: A.K., R.T., D.N.; resources: A.K., D.N., T.G., B.M.R., J.C.K.; data curation: R.T., D.N.; original draft preparation: A.K., R.T.; review and editing: A.K., R.T., B.M.R., J.C.K.; supervision: A.K.; project administration: A.K.; funding acquisition: A.K., T.C.G., B.M.R. and J.K. All authors have read and agreed to the published version of the manuscript.

